# A lipidomic exploration of the effects of high-intensity interval exercise in healthy men after metformin intake

**DOI:** 10.1101/2025.04.02.646809

**Authors:** René Neuhaus, Thomas Meikopoulos, Ana Gradillas, Carolina González-Riaño, Georgios Theodoridis, Stefanos Nikolaidis, Vassilis Mougios, Coral Barbas, Helen Gika, Alma Villaseñor

## Abstract

We have previously found that high-intensity interval exercise (HIIE) affected metformin pharmacokinetics, causing higher maximal plasma concentration compared with rest. In this scenario, changes in individual lipids could play an important role. The prolonged responses of the lipidome to HIIE have not been explored.

Thus, this study aimed to explore differences in the plasma lipidomic profiles between HIIE and rest, both under metformin treatment.

Nine healthy males participated in two sessions where they received 1,000 mg of metformin. In session A, they performed HIIE at an average intensity of 67% of maximum heart rate for a total duration of 76 min, whereas in session B they rested. Plasma was collected before taking metformin and during each session (in total 14 time points spanning 12 h). Samples were analysed through lipidomics using mass spectrometry. Paired Wilcoxon tests between sessions were applied for statistics.

We found several variations in the lipid profiles due to HIIE, which persisted until 4 h post-exercise. The main discriminant lipid classes were fatty acids, acyl carnitines, glycerophosphocholins, sphingomyelins, and triglycerides. These changes were followed in time up to 12 h, showing the effect of the meals taken during the session. We hypothesize the changes are a synergic effect of HIIE and metformin in the lipidome with the effect of HIIE being the predominant.

These findings provide important insights into the dynamic and complex physiological response of humans to intensive exercise under metformin intake.

## INTRODUCTION

Maintaining adequate levels of physical activity is essential for good health. Physical activity is widely recommended to prevent and treat obesity-related complications such as type 2 diabetes mellitus (T2DM)(1). Recent studies have shown that high-intensity interval exercise (HIIE) which includes alternating short periods of intense exercise with brief recovery periods, is effective in improving cardiometabolic markers in patients with insulin resistance and T2DM(2–4). Physical exercise is known to produce large changes in the concentrations of multiple metabolites(5), including lipids(6). Among the latter, ceramides (Cer), diglycerides (DG), and acyl carnitines (CAR) have been shown to activate intracellular signalling cascades and regulate metabolic pathways(7–11). It has been described that physical exercise induces lipid alterations such as an increased utilization of triglycerides (TG) that in exchange increase the concentration of fatty acids (FA) and CAR(12). Other studies have addressed the regulation of the TG in the skeletal muscle upon exercise(13,14), The overall regulation of circulating lipid species to HIIE is poorly understood. To cope with this challenge, lipidomic analysis employing high throughput techniques, such as liquid chromatography coupled to mass spectrometry (LC-MS) can be used, as it allows the acquisition of a lipid profile consisting of numerous lipid species(2–4).

Metformin is a first line antihyperglycemic drug for T2DM treatment(15). As a first step in the exploration of the relationship between metformin administration and HIIE, we have previously shown that HIIE during the absorption phase of metformin affected metformin pharmacokinetics, causing higher maximal plasma concentration (C_max_) compared with rest without compromising glucose homeostasis in healthy men(16). In this scenario, changes in the plasma concentrations of individual lipids could play an important role in the response to HIIE. However, the way that the plasma lipidome changes, over consecutive hours in a controlled environment of food intake and either HIIE or rest, has not been explored.

Thus, the aim of this work was to explore the changes in the plasma lipidome of healthy individuals for 12 h after metformin intake followed by either HIIE or rest.

To our knowledge this is the first study reporting HIIE effect on the blood lipidome of healthy individuals over a long period of time covering exercise session and post-exercise, using a comprehensive, untargeted lipidomic approach.

## RESULTS

To study the effect of exercise on the lipidome, we collected plasma samples from 9 healthy men that participated in two sessions that lasted 12 h, one including 76 min of HIIE, the other one resting **(Figure 1)**. In both sessions the participants took 1,000 mg of metformin at the beginning. For practical reasons, from all collected samples, a subgroup was analyzed via reverse phase ultra high-performance liquid chromatography coupled to mass spectrometry with an electrospray ionization source and a quadrupole–time of flight mass analyzer (RP-UHPLC-ESI-QTOF-MS).

**Figure 1.**
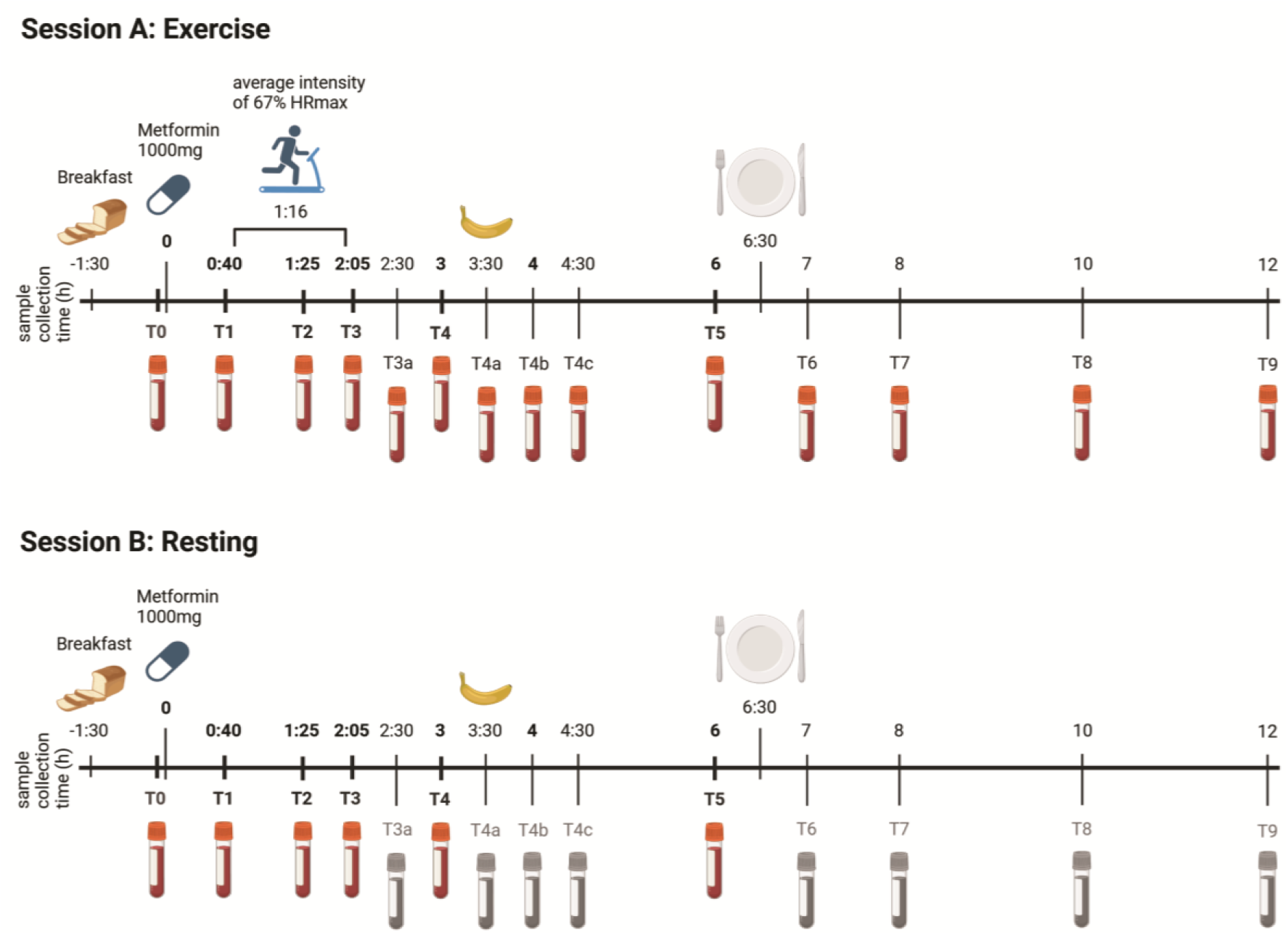
Study design. Outline of the two sessions that participated the 9 healthy subjects showing food intake, metformin intake, blood sampling, and exercise. Blood samples in red means samples were analysed using lipidomics (n=9 in each time point).

The samples collected at the same time point in both sessions (T0-T5) were analyzed and compared (**Figure S1**). The untargeted lipidomic analysis yielded a total of 1,303 and 299 chemical features in ESI(+) and ESI(–) modes, respectively. The unsupervised model of these features using PCA showed tight clustering of QC samples in the plots for each ionization mode (**Figure S2A, B**). After lipid annotation from both ionization modes and removal of duplicates, a single data matrix was obtained consisting of 247 unique lipids (**Table S1**). The PCA model of these data continued to show clustering of QC samples (**Figure S2C**) indicating good quality of the data and that the variation shown is due to inter-individual variation of lipids in the plasma of the participants. Interestingly, PCA models based on the samples alone from all-time points did not reveal any discrimination between rest and exercise sessions (**Figure S2D-F**). Thus, further statistical analysis was applied to get a deeper insight into the data. No sample outliers were detected with the profile of 247 annotated lipids (**Figure S2F**).

### Lipidomic changes after HIIE involve a complex network of different lipid classes

To study the effects of HIIE on the plasma lipidome, the time points T0-T5 were considered separately and lipidome was compared under rest and exercise conditions using the Wilcoxon test. Specific focus was set on a time frame up to 6 hours (**Figure S1**), where early effects might be reflected by HIIE. Univariate statistical analysis showed several lipid differentiations (**Figure 2A**, **Tables S2-S5**).

**Figure 2.**
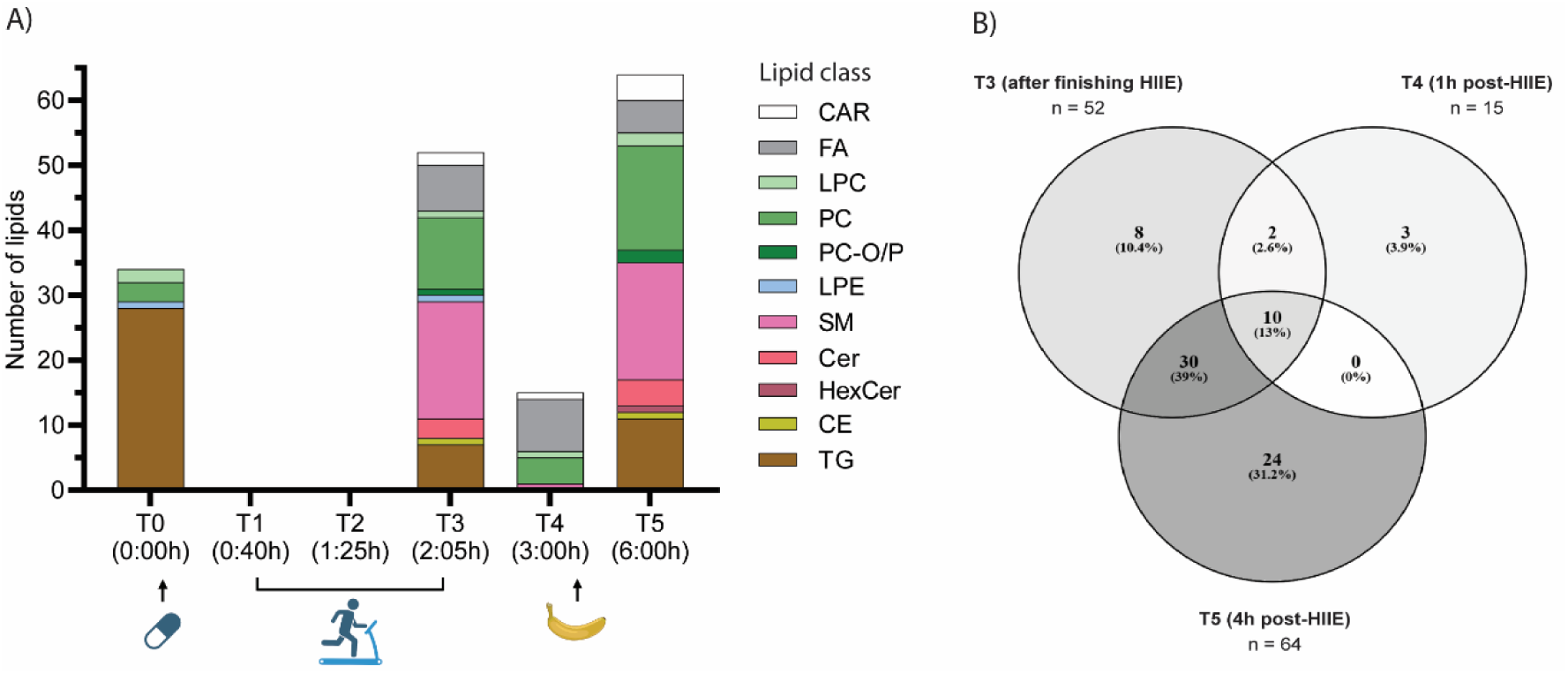
Significantly different lipid species between exercise and resting session under metformin intake. **A)** Stack bar plot of significantly different lipids (p < 0.05, pFDR< 0.2) per lipid species at different time points after Wilcoxon test between exercise and resting sessions for the 9 healthy subjects taking metformin. Abbreviations: acyl carnitines (CAR), fatty acids (FA), lysoglycerophosphocholines (LPC), glycerol-phosphocholines (PC), ether-glycerophosphocholines (PC-O/P), lysoglycerophosphoethanolamines (LPE), sphingomyelins (SM), ceramides (Cer), hexosylceramides (HexCer), cholesteryl esters (CE), triglycerides (TG). **B)** Venn diagram showing shared significant lipids after Wilcoxon test between exercise and resting sessions for the 9 healthy subjects taking metformin at T3 (after finishing HIIE), T4 (1 h post-HIIE) and T5 (4 h post-HIIE). The intensity of the grey color corresponds to the amount of shared lipids. Abbreviations: high-intensity interval exercise (HIIE).

Starting with T0 (1.5 h after breakfast) it was found that 34 lipids had significantly different levels between the two sessions (p < 0.05, pFDR < 0.2). These lipids belong to TG, glycerophosphocholines (PC), lysoglycerophosphocholines (LPC), and lysoglycerophosphoethanolamines (LPE). Their connections were presented using lipid networks by Linex as shown in **Figure S3A**. As can be seen, most of the TGs, PCs, and LPCs were connected within their classes, with TGs (e.g. TG(44:1), TG(42:1), and TG(42:0) exhibiting the highest fold change in exercise vs rest. All these lipids were higher in the exercise session before HIIE mostly probable due to a food intake effect (**Table S2**).

These lipid differences found at T0 seem to fade out at T1 and T2, which correspond to prior and 40 minutes into exercise, respectively. Interestingly, it was after 76 minutes of HIIE, at T3, that 52 lipid species (**Table S3**) emerged as statistically different from resting conditions (p < 0.05, pFDR < 0.2). The main lipid classes of these species were sphingomyelins (SM), PC, TG, and FA, followed by CAR and Cer (**Figure 2A**). The lipid network suggested again that most of the lipids were connected within their classes and that acetylcarnitine [CAR(2:0)] and palmitoleic acid [FA(16:1)] were the lipids with the highest fold change **(Figure 3A**). Almost in all cases, except for LPE(22:6), increased levels of these lipids were observed in the plasma of the exercise group (**Table S3**).

**Figure 3.**
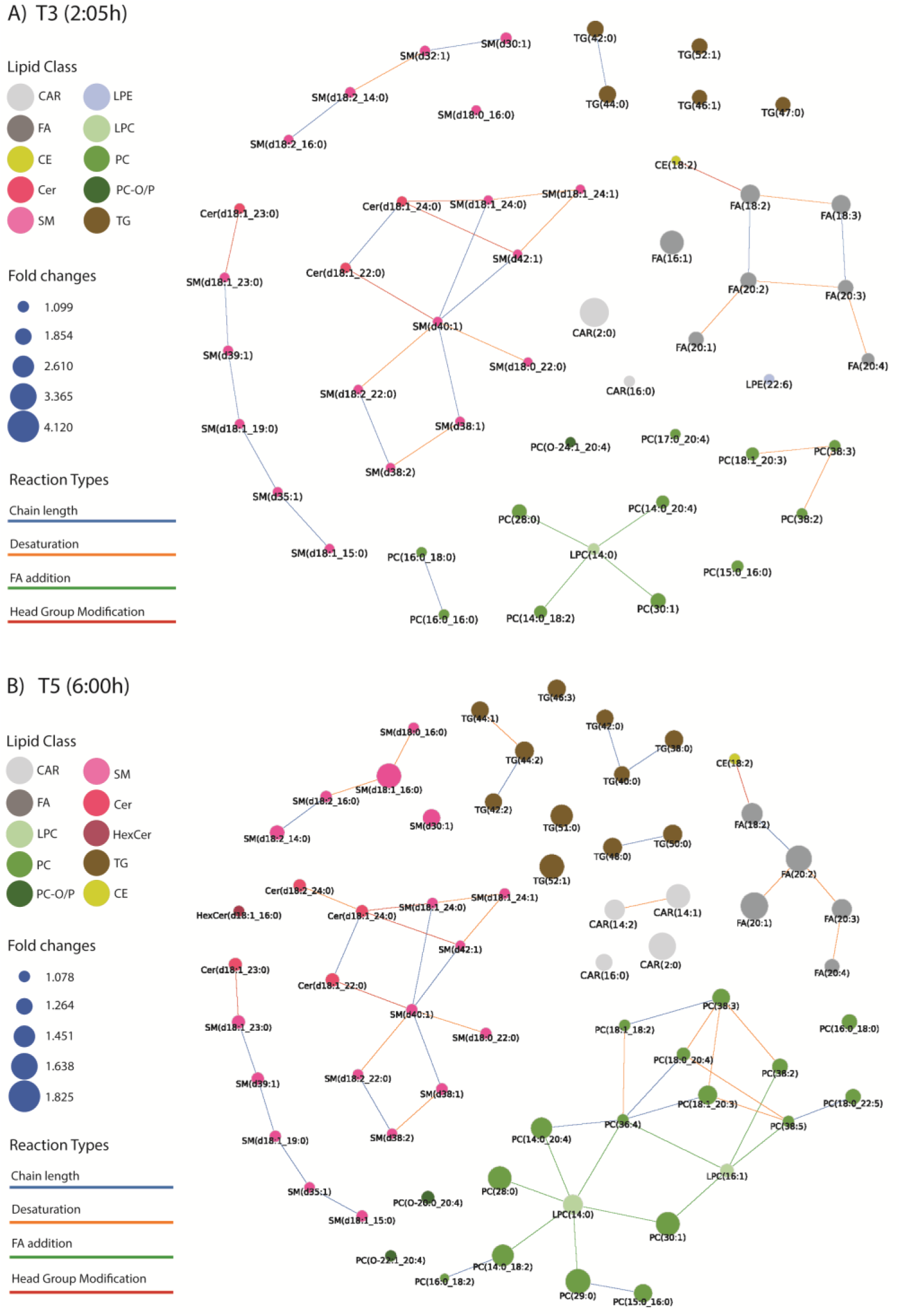
Lipid networks of the statistically significant lipids at T3 and T5. Lipid network at **A)** T3 (2:05 h) just after finishing HIIE and **B)** T5 (6:00 h) 4 hours after finishing HIIE and after a snack between exercise and resting sessions for the 9 healthy subjects taking metformin. Each dot represents a lipid, the size represents the fold change from individuals at exercise session against resting conditions. The color of the dots represents the lipid class. All lipid names are depicted as the composition (“_”), to see the *sn-*positions (“/”) see Tables S3 and S5. Lipids with the same sum composition are represented as one lipid. Abbreviations: acyl carnitines (CAR), fatty acids (FA), lysoglycerophospho-cholines (LPC), glycerophosphocholines (PC), ether-glycerophospho-cholines (PC-O/P), lysoglycero-phosphoethanolamines (LPE), sphingomyelins (SM), ceramides (Cer), hexosylceramides (HexCer), cholesteryl esters (CE), triglycerides (TG).

After almost one hour of finishing HIIE at T4 only 15 significant lipid species were significantly different, in almost all cases increased with the exception of LPC(16:0), compared with resting conditions (p < 0.05, pFDR < 0.2). These included CAR(2:0) and members of the FA and PC classes (**Figure S3B**, **Table S4**). The lipid network shown in **Figure S3B** revealed the importance of CAR(2:0) and FAs such as, FA(20:1), FA(20:2), and FA(18:2). Moving away from the completion of the exercise session, three hours later (at T5, six hours from beginning of the session and after banana intake), 64 lipids were found increased in the exercise group (**Table S5**). Again, the lipid classes that mainly found differentiated were PC, SM, and TG **(Figure 2A).** Lipid network pointed out additional lipid classes, such as FA and CAR, which showed high fold changes (**Figure 3B**). A Venn diagram comparing the significant lipid changes between T3 (after finishing HIIE), T4 (1 h post-HIIE) and T5 (4 h post-HIIE) is shown in **Figure 2B**. In this figure, we observed that T3 and T5 shared around 50% of the significant lipids showing the strong effect of HIIE on the lipidome even after 4 h post-exercise.

To understand better the changes observed, the levels of some of the most altered lipid species due to exercise such as CAR(2:0), FA(16:1), FA(20:1), FA(20:2), FA(20:3), and FA(20:4) were plotted for the two sessions (**Figure 4**). It can be observed that CAR(2:0) increased dramatically after starting HIIE (T2) and continued increasing, reaching its highest level right after finishing exercise (T3). One hour afterwards (T4), a decrease was observed, nevertheless, even 4h after the termination of HIIE (T5) the level is still higher than in resting conditions.

**Figure 4.**
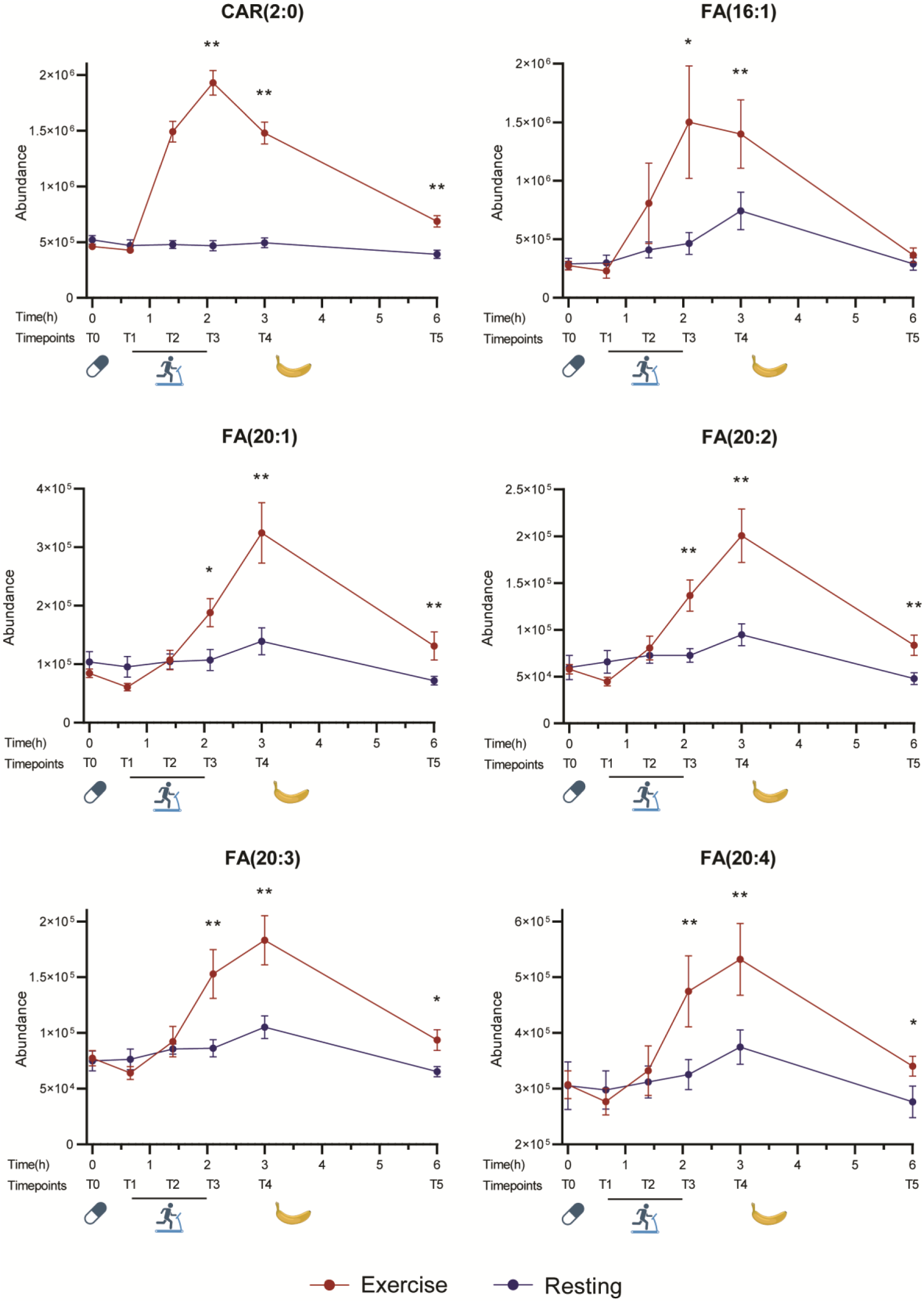
Lipid trajectories for lipids with high fold changes due to HIIE from T0 (0 h) to T5 (6 h). Abundances in exercise and resting sessions of CAR(2:0), FA(16:1), FA(20:1), FA(20:2), FA(20:3) and FA(20:4). Values are represented as mean of normalized area ± SEM of 9 healthy subjects taking metformin, significant differences were found using Wilcoxon test; * p< 0.05; ** p< 0.01 between exercise and resting sessions.

In the case of the FA, a rise in their level was also observed, which however continued to increase after the completion of HIIE, reaching a maximum at T4, 1 h after HIIE. Similarly to CAR(2:0), most FAs did not return to the levels at resting conditions at T5.

Other cases of altered lipids, such as PC(16:0_18:0), LPE(22:6), SM(d40:1), Cer(d18:1/22:0), and some TGs; TG(56:2), and TG(42:0), exhibited diverse trajectories indicating mainly an increased effect around the exercise time window (T1-T3) that tends to get back at rest levels later on (**Figure S4**). PC(16:0_18:0), SM(d40:1), Cer(d18:1/22:0), TG(56:2), and TG(42:0) increased upon HIIE (to a lower extent compared to CAR and FA) whereas LPE(22:6) showed a decreasing trend.

### Trajectories of exercise-induced lipid changes over 12 hours show interaction with meals

To further examine the trend of lipids most affected by exercise beyond 6 h (T5), all 14 sampling points of the exercise session, up to 12 h (T9), were plotted (**Figure 5**). These graphs show that CAR(2:0) continued decreasing, reaching baseline (that is, the pre-exercise values) at 12 h, regardless of the food intake in between. In the case of the FA, the downward trajectory starting after the completion of exercise already described above, was interrupted by the meal taken at 6.5 h, after which the FA concentrations dropped again (**Figure 5**). This pattern was more pronounced with FA(16:1) but less with FA(20:1), FA(20:2), FA(20:3), and FA(20:4).

**Figure 5.**
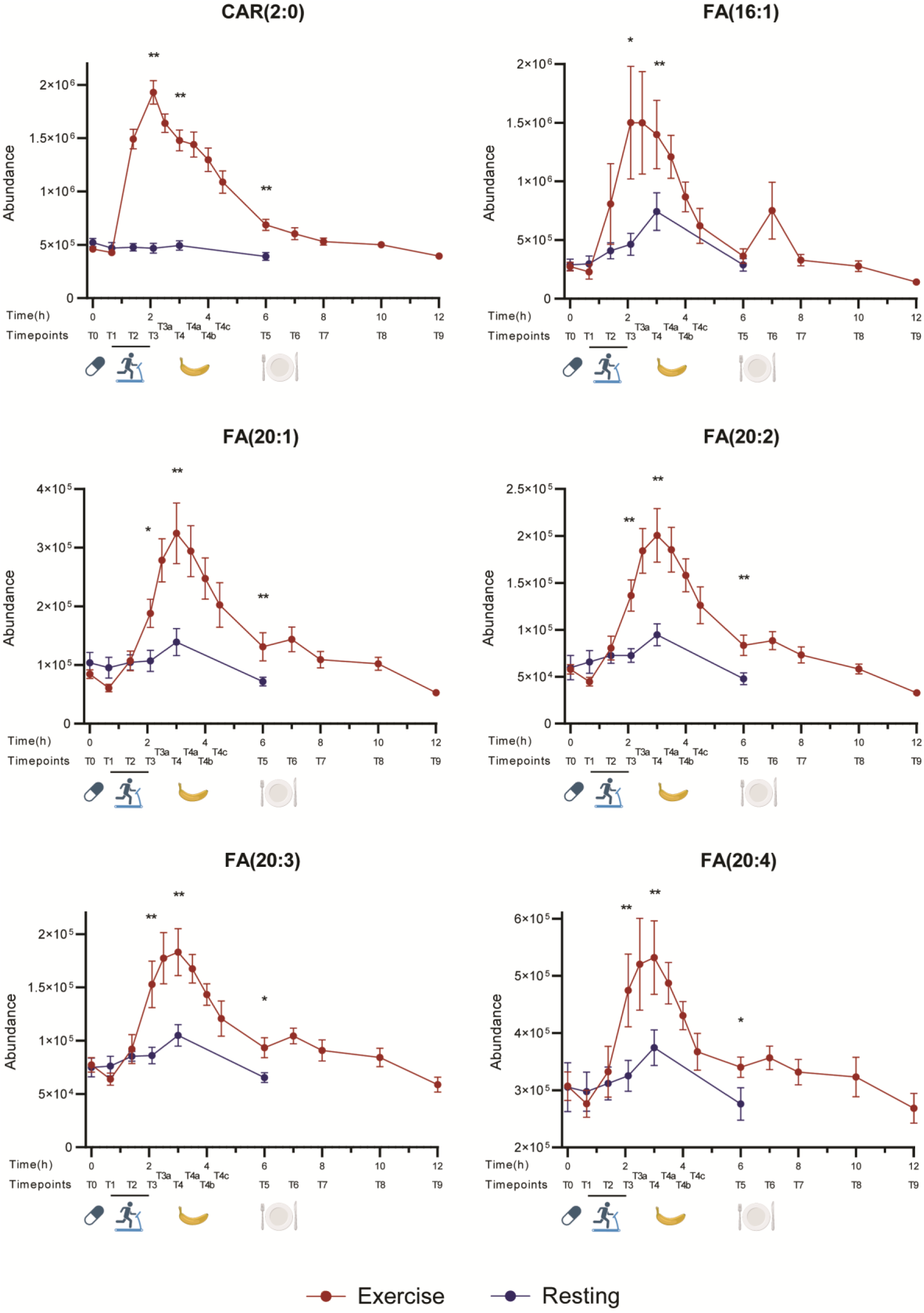
Lipid trajectories for lipids with high fold changes due to HIIE from T0 (0 h) to T9 (12 h). Abundances in exercise and resting sessions of CAR(2:0), FA(16:1), FA(20:1), FA(20:2), FA(20:3) and FA(20:4). Values are represented as mean of normalized area ± SEM (n=9 healthy subjects taking metformin), significant differences were found using Wilcoxon test; * p< 0.05; ** p< 0.01 between exercise and resting sessions.

The 12 h trajectories of the lipid species that were affected less by exercise are presented in **Figure S5**. In these cases of lipids, like PC(16:0_18:0), LPE(22:6), SM(d40:1), and Cer(d18:1/22:0) the levels after 6 h did not change considerably and no impact of food intake was detected. Regarding the altered TGs, it was found that TG(56:2) strongly increased after the meal, peaking at 10 h, whereas TG(42:0) increased to a lower extent and only transiently after the meal, after which it continued dropping.

Overall, the trends of all annotated lipids in relation to exercise can be seen in the heatmap which shows the average intensities at each of the 14 time points from 0 to 12 h (**Figure 6**). It can be clearly seen that plasma CAR, FA, SM, some specific TG (saturated and unsaturated), and PC increased during and after HIIE (T2 and T3), whereas LPC decreased. From the increased lipids, TG and PC decreased 30 min post-exercise, whereas CAR and FA remained elevated until 2 hours after HIIE (T4b-T4c). It can be also noted that, after the meal (T6 and on), mainly PC, LPC, and TG (especially polyunsaturated TG) increased.

**Figure 6.**
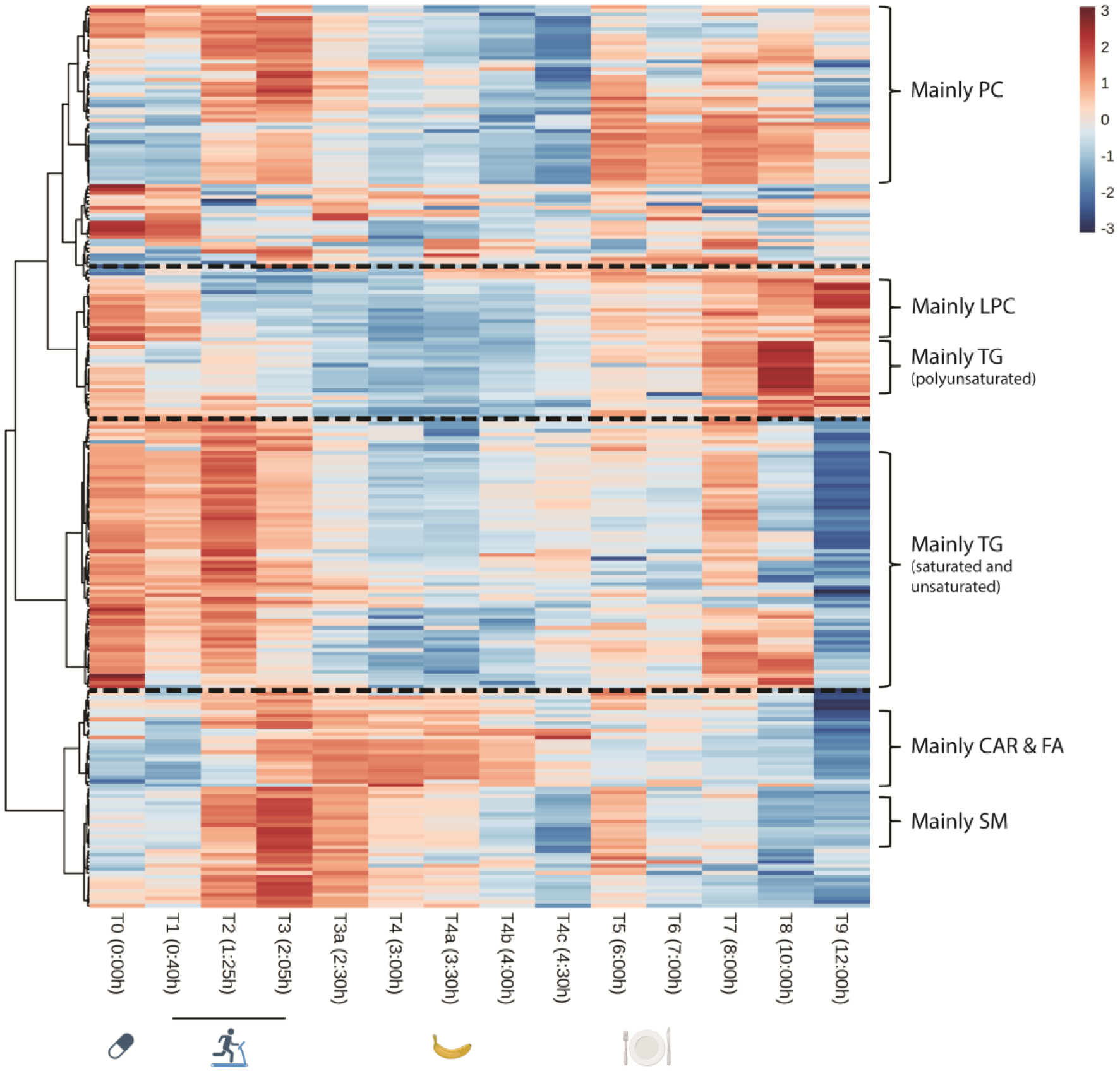
Trends of lipid classes. Descriptive diagram. Heatmap of average abundances of all annotated lipids at all time points (12 h) in exercise session. Abbreviations: acyl carnitines (CAR), fatty acids (FA), sphingomyelins (SM), triglycerides (TG), lysoglycerophosphocholines (LPC), glycerophosphocholines (PC). Each time point (T0-T9) shows the average of the 9 healthy subjects taking metformin during the exercise session.

## DISCUSSION

It has been reported that HIIE interventions may play a role in improving long-term cardiovascular health(17) via favourable changes in the blood lipid content such as decreasing low-density lipoprotein (LDL) and total cholesterol (TC) and increasing high-density lipoprotein (HDL), following high-intensity exercise(18). Increased exercise intensity correlates with high levels of catecholamines, cortisol, and growth hormone which have a lipolytic effect and in combination to the additional increase in fat oxidation that occurs, it can positively affect levels of circulating lipids(19). The lipolytic effect occurs as breakdown of TG in adipose tissue releases free fatty acids (FFA) into the bloodstream. FFA serve as an energy source for muscles and other tissues but their oxidation is suppressed during high-intensity exercise (above ∼75% VO₂ max) as a shift toward carbohydrate metabolism occurs(20). In parallel, muscle contraction increases lipoprotein lipase (LPL) activity, which leads to declined levels of circulating TG(21).

Even a single HIIE session can lead to immediate TG reduction, temporary increase in HDL, and sustained fat oxidation post-exercise(22–25). Increased fat oxidation is sustained post-HIIE and it is shown that FFA remain elevated for a period as the body continues to rely on fat metabolism for recovery and decreased levels of TG may persist for hours after exercise due to enhanced clearance by LPL(26). However, the findings are diverse as the effect on lipid metabolism is related to the intensity of the exercise and the variations in the HIIE protocol applied.

Up to now, scarce data exist on the effect of HIIE by comprehensive lipidomic analysis. Studies are very limited on the effect of HIIE on the plasma lipidome of healthy individuals and have been performed only for comparative purposes to other type of exercise after repetitive sessions over a period of time(27), either in combination with diet(28) or in comparison to a disease group(3). In most of the cases, studies performed of the effect of HIIE present data limited to typical blood lipid parameters like cholesterol, LDL, HDL and TG(29,30). In these cases, positive impact on lipid metabolism has been shown. However, there is no further evidence on the effect of HIIE on diverse lipid species. In this study, by performing untargeted lipidomic analysis we were able to capture information for 247 unique lipid species from various classes.

Our findings show that significant changes occur in different lipid species during and after HIIE, suggesting complex correlations networks. CAR and FA species showed to be mostly affected with a steep increase in their levels, while other classes of lipids such as SM, PC, LPE, Cer and TG also showed alterations.

The applied methodology captures comprehensively the impact of HIIE on the plasma lipidome, thus changes in lipid metabolism during and after HIIE over a period of 12 h in healthy individuals are shown for the first time.

In one of the very few relevant lipidomic studies where a wide range of lipid classes was assessed in serum with the aim to compare a 6-week moderate-intensity continuous training group with sprint interval training(31), alterations in various classes of lipids have also been observed. SM(d18:1/22:5), SM(d18:1/22:6), MG(18:2), LPE(18:1), LPE(20:1), PC(O-16:0/16:0), and PI(16:1/18:1) were found increased, while most PC, Cer and hexosylceramides (HexCer) were decreased. Certainly, that study focuses on different training protocols, however, its findings via a holistic approach indicate the diversity of lipids altered as a result of exercise.

In our study we have observed acute increase of acetyl carnitine, CAR(2:0), which has been reported in several studies, suggesting that exercise leads to increased carnitine release into the plasma, especially when glycogen stores are depleted, as the body shifts more towards fat metabolism. The rapid increase in acetyl-CoA production and the role of carnitine as acetyl buffering in skeletal muscle metabolism led to CAR(2:0) build up in muscle which then is released into the bloodstream(32).

Significant FA rise observed in the study is also in agreement with lipolysis activation and break down of stored TGs expected to occur in the training session(33). SM and ceramides found elevated are known to be involved in stress signalling and muscle adaptation, whereas PC are major component of cell membranes that may play a role in lipid remodeling to maintain membrane integrity during HIIE and muscle cells stress(34,35). Lipid mobilization and sharp increase in energy demand during HIIE can also explain the observed increase of some TG. These observations are the reflection of complex metabolic perturbations due to a strong factor such as intense exercise, while metformin intake should not be disregarded.

Although earlier, based on studies measuring basic lipid characteristics e.g. TG, cholesterol and HDL, LDL, it was considered that acute metformin intake did not affect circulating lipid concentrations(36), more recent studies applying comprehensive lipidomic analysis approaches have indicated that metformin can alter plasma lipid composition, including decrease in saturated fatty acids, associated with insulin resistance, increase in polyunsaturated fatty acids and altered ceramide levels. Dahabiyeh *et al*(37) have recently provided evidence by applying a mass spectrometry shotgun lipidomic approach that a single dose of 500 mg metformin induced acute alterations in the plasma lipidome of healthy subjects. Thirty-three lipids were significantly increased and 192 decreased in plasma in response to acute metformin intake. The altered lipids are mainly involved in arachidonic acid metabolism, steroid hormone biosynthesis, and glycerophospholipid metabolism. It was also shown that metformin had the most impactful effect on lipidome at its Cmax. Furthermore, several lipids displayed a similar change in their levels compared to metformin, among which are monoacylglycerols; a larger number of lipids acted in an opposed manner to metformin levels, including fatty acyls (fatty esters, fatty acids, and eicosanoids), sterol lipids, glycerolipids, and glycerophospholipids (PC, glycerophosphoinositols)(37). In an earlier study from the same group(38), several lipids were revealed with opposed trends to metformin patterns, mainly involving arachidonic and linoleic acid metabolism and including glycerophospholipids (such as lysoglycerophosphates (LPA) and LPC). It was concluded that the pleiotropic effect of metformin probably is associated with its effect in lipid metabolism and lipid signalling pathways(38). Hence, we could hypothesize that, although to a smaller extent than exercise, metformin intake could have played a role on the lipid dysregulation observed in this study and that a combinatorial effect in lipid metabolism might have been captured with the HIIE being the predominant. Taking into the account that in one of our previous studies in healthy male individuals, we observed that the levels of metformin in plasma were higher during HIIE compared to resting conditions(16), an increased impact might be expected.

It could also be mentioned that a similar trend with that described by Dahabiyeh *et al*(37) regarding the similar trend of altered lipids with the metformin levels along time have been noted in our study. CAR(2:0) maxima coincided with that of metformin and they seem to follow similar trajectories over time (**Figure S6A**). A similar curve trend of CAR found in our study was also noted by Dahabiyeh *et al*(37). On the other side, TG(56:2) shows an opposite trend to that of metformin overtime (**Figure S6B**).

These findings could be of high relevance for pathologies such T2DM, where metformin is administered and routine exercise is recommended. Recently HIIE has been popularized among nonathletes for improving cardiovascular health(39), weight loss(40), and insulin sensitivity(41). Providing deeper insight on the biochemical modulations triggered by high intensity exercise could provide further knowledge to enhance its beneficial impact in health, as the complex metabolic effects that are induced are yet not fully understood.

Definitely, further investigation is required to enlighten the biochemical processes behind the current findings and to evaluate their importance to improve the contribution of intensive exercise in the battle against metabolic diseases such as obesity and diabetes. To our knowledge, this is the first study that monitored the effect of HIIE on the plasma lipidome under metformin for a period of 12 h. A limitation of the study is the small number of participants, which was primarily due to its complex and demanding design. Additionally, as based on previous evidence, acute intake of a therapeutic dose of metformin had probably modest effect on plasma lipids, therefore the interaction between metformin and HIIE effect is difficult to interpret. Further studies with larger sample sizes and a control group without metformin intake are needed to confirm or refute our findings.

All in all, this study helps to understand better the highly dynamic lipid alterations in blood by HIIE after metformin intake. The main discriminant lipid classes were fatty acids, acyl carnitines, glycerophosphocholins, sphingomyelins, and triglycerides. These lipid classes continued to be significant up to 4 h after finishing HIIE.

## MATERIALS AND METHODS

### Study design

This work is part of a wider study examining the effect of exercise on the pharmacokinetics of metformin(16). In our study, 9 healthy, non-smoking male subjects participated in two sessions, A and B, each lasting 12 h, performed in random counterbalanced order with one or two weeks in between.

*Inclusion and exclusion criteria:* Participants were non-smoking males, who were not taking any medication and abstained from drinking alcohol and coffee for 72 and 12 h, respectively, before each experimental session, as well as during the sessions. Participants were informed about the purpose, benefits and risks of the study and then gave their written consent to participate. The study was in accordance with the Helsinki Declaration and was approved by the Research Ethics Committee of the School of Physical Education and Sport Science at Thessaloniki, Aristotle University of Thessaloniki, following the Regulation of Principles and Operation of the Research Ethics Committee of the Aristotle University of Thessaloniki and the applicable legislation (approval number 113/2022).

In both sessions, the participants received 1,000 mg of metformin (Glucophage^®^, immediate-release tablets) at around 10 am, 1-1.5 hours after receiving a light, low-fat, high-carbohydrate breakfast. In session A, they performed a HIIE protocol with an average intensity of 67% of maximum heart rate (HRmax) and a total duration of 76 min. HIIE was performed 45 min after taking metformin and consisted of 5 min of warm-up by walking at 4-5 km/h, followed by 11 sets of 6 min, each consisting of 3 min of walking or running at 90% of HRmax and 3 min of walking at 4-5 km/h. Between sets 6 and 7, a 5-min rest was included. The participants then rested throughout the remainder of the day. In session B, they rested throughout the whole day.

In both sessions, the participants consumed one or two bananas at 3.5 h after taking metformin and a regular serving of spaghetti at 6.5 h. Schematically, the study design is presented in **Figure 1**.

### Sample collection and processing

Fourteen venous blood samples were collected in EDTA tubes in each session at the Laboratory of Evaluation of Human Biological Performance as follows: before taking metformin (0 h, T0) and after taking metformin at 40 min (T1), 1 h 25 min (T2), 2 h 5 min (T3), 2 h 30 min (T3a), 3 h (T4), 3 h 30 min (T4a), 4 h (T4b), 4 h 30 min (T4c), 6 h (T5), 7 h (T6), 8 h (T7), 10 h (T8) and 12 h (T9) (**Figure 1**). Plasma was collected by centrifugation and stored at –80 °C until shipment to CEMBIO, where it was kept at –80 °C until analysis. For practical reasons, only samples taken at T0, T1, T2, T3, T4, and T5 of session B were subjected to lipidomic analysis, whereas all samples taken during session A were analyzed. Thus, leading a total of 180 individual samples to be analyzed.

### Reagents and chemicals

For all aqueous solutions, reverse-osmosed ultrapure water was obtained from a Milli-Qplus185 system (Millipore, Billerica, MA, USA). LC-MS-grade isopropanol, methanol, and acetonitrile were purchased from Fisher Scientific (Pittsburgh, PA, USA). Methyl *tert-*butyl ether (MTBE) (HPLC grade, ≥ 99.8%), ammonium fluoride (ACS reagent, ≥ 98%), and palmitic acid-d31 were purchased from Sigma-Aldrich (Steinheim, Germany). Analytical grade ammonia solution (28%, GPR RECTAPUR^®^) and glacial acetic acid (AnalaR® NORMAPUR^®^) were obtained from VWR Chemicals (Radnor, PA, USA). Sphinganine (d17:0) primary standard and LightSPLASH^®^ LIPIDOMIX^®^ Quantitative Mass Spec Primary Standard were obtained from Avanti Polar Lipids (Alabaster, AL, USA).

### Sample preparation for lipidomics

Lipidomics analysis was carried out following previously published methodology (42). Plasma samples were thawed on ice for approximately 1 h and vortex-mixed for 2 min. Then, 50 μL of each sample were transferred to an Eppendorf vial on ice. Subsequently, 175 μL of ice-cold methanol (stored at –20 °C), containing 2.3 ppm of sphinganine (d17:0) and 4.6 ppm of palmitic acid-d31 as internal standards, were added for deproteinization. After shaking the sample for 1 min, 175 μL of MTBE and 10 μL of LightSPLASH LIPIDOMIX Quantitative Mass Spec Primary Standard mix were added. Samples were vortex-mixed for 30 min and centrifuged for 15 min at 15 °C and 16,100 x *g*. Finally, 100 μL of supernatants were added to chromatography vials, equipped with 300-μL inserts (one for each ionization mode, positive and negative), centrifuged for 5 min at 15 °C and 2,000 x *g*, and then injected into the LC-MS system (**Figure S7A**).

In addition, 7 types of quality control samples (QC) were prepared to control the quality of the analysis and to facilitate lipid annotations. First, a “QCpool” sample was prepared by mixing 5 μL of each of the 180 analyzed samples. Then, the rest of the QC samples were used for iterative MS/MS experiments, a “QC0 (A+B)” sample was prepared by mixing 15 μL of samples taken at T0 of both sessions (18 in total). Additionally, a “QC1 (A)” sample was prepared by mixing 25μL of samples collected at T1 of session A (9 in total). Similarly, “QC5 (A)” (T5 of session A), “QC1 (B)” (T1 of session B), “QC5 (B)” (T5 of session B), and “QC9 (B)” (T9 of session B) samples were prepared.

### Analysis in Liquid chromatography coupled to QTOF mass spectrometry analysis (LC-QTOF-MS)

Samples were analyzed using an Agilent 1290 Infinity II UHPLC system coupled to an Agilent 6545 quadrupole time-of-flight (QTOF) mass spectrometer (Agilent, Santa Clara, CA, USA). An Agilent InfinityLab Poroshell 120 EC-C18 (3.0 × 100 mm, 2.7 µm) column, protected by a compatible guard column (Agilent InfinityLab Poroshell 120 EC-C18, 3.0 × 5 mm, 2.7 µm), was employed for the analysis. The column temperature was set at 50°C, while the autosampler’s temperature was set at 15 °C to preserve compound stability and prevent lipid precipitation. Data were acquired in both ionization modes in separate runs. For positive ionization (ESI+), the injection volume was set at 0.5 μL, while, for negative ionization (ESI-), at 1 μL. Elution was performed with a solvent system consisting of (A) 10 mM ammonium acetate, 0.2 mM ammonium fluoride in water/methanol (9:1, v/v) and (B) 10 mM ammonium acetate, 0.2 mM ammonium fluoride in acetonitrile/methanol/isopropanol (2:3:5, v/v/v). The elution program applied was 70% B at 0–1 min, linear gradient to 86% B until 3.5 min, held until 10 min, linear gradient to 100% B until 11 min, and held until 17 min. Then, the system returned to the initial condition to achieve column equilibration for a total of 2 min. Thus, the total analysis time was 19 min.

The QTOF mass spectrometer, equipped with a dual atmospheric jet stream electrospray ionization (ESI) ion source, was configured with the following parameters: 150 V fragmentor, 65 V skimmer, 3500 V capillary voltage, 750 V octopole radio frequency voltage, 10 L/min nebulizer gas flow, 200 °C gas temperature, 50 psi nebulizer gas pressure, 12 L/min sheath gas flow, and 300 °C sheath gas temperature. In full scan mode, operated from 50 to 1.800 *m/z* with a scan rate of 3 spectra/s, data were collected for both ionization modes. A solution of two reference mass compounds was used throughout the analysis: purine (C_5_H_4_N_4_) at *m/z* 121.0509 for the positive and *m/z* 119.0363 for the negative ionization modes; and HP-0921 (C_18_H_18_O_6_N_3_P_3_F_24_) at *m/z* 922.0098 for the positive and *m/z* 980.0163 (HP-0921 + acetate) for the negative ionization modes. These masses were continuously infused into the system to provide constant mass correction.

At the end of the analytical run, the 7 types of QC were used to perform iterative runs of tandem mass spectrometry (MS/MS, 5 for each type) in both ionization modes. They were analyzed with MS and MS/MS scan rates of 3 spectra/s, 40-1,700 *m/z* mass window, narrow (∼ 1.3 amu) MS/MS isolation width, 3 precursors per cycle, 5,000 counts, and 0.001% of MS/MS threshold. Five iterative-MS/MS runs for each QC sample were set at a collision energy of 20 eV, and the subsequent five runs for each QC sample were performed at 40 eV. To avoid inclusion in the iterative-MS/MS data, reference masses and contaminants detected in blank samples were excluded from the analysis.

### Data treatment

The Agilent MassHunter Profinder B.10.0.2 software was used to reprocess the data collected after the LC-MS analyses in both positive and negative ion modes, yielding 1,303 and 299 features, respectively. The datasets were extracted using the Batch Recursive Feature Extraction (RFE) algorithm integrated into the software. The selected adducts were the following: [M+H]^+^, [M+Na]^+^, [M+K]^+^, and [M+NH_4_]^+^ in ESI(+); [M-H]^−^, [M+CH_3_COO]^−^, and [M+Cl]^−^ in ESI(–). The neutral loss of water was also considered for both ion modes.

Samples were analyzed only once in the instrument in both ionization modes leading to one analytical replicate. The quality of their analysis was assured by the accomplish of the quality assurance (QA) strategy based on data filtration and normalization. Missing values were estimated using the *k-*Nearest Neighbor algorithm(43) in MATLAB (2018, MathWorks, Netik, MA, USA). Then, the features were filtered based on the coefficient of variation of the QC samples, with a cutoff threshold of 30%. As the number of samples was high, data were normalized using the quality control samples and support vector regression (QC-SVRC) algorithm(44) in MATLAB. Finally, to ensure the quality of the obtained data and detect sample patterns and possible outliers, we applied principal component analysis (PCA) models using SIMCA 16 (Sartorius, Göttingen, Germany).

### Lipid annotation

The workflow for lipid annotation consisted of three steps (**Figure S7B**): First, the raw LC-MS/MS data obtained were imported into the Lipid Annotator software (Agilent). The software created a fragmentation-based MS/MS library containing the *m/z* and matching retention time (RT) of all precursors recognized as lipids. The Lipid Annotator method was set as follows: ion species [M+H]^+^, [M+Na]^+^, and [M+NH_4_]^+^ for ESI(+); and [M-H]^−^ and [M+CH_3_COO]^−^ for ESI(–). Then, for both ion modes, the Q-score was set at ≥ 50; all lipid classes were selected, mass deviation was established as ≤ 20 ppm, fragment score threshold was fixed as ≥ 30, and total score was set at ≥ 60.

Second, raw LC-MS/MS data were reimported into the open-source MS-DIAL software(45) in order to achieve an additional lipid identification with the aim of combining the information from both software to confirm lipid annotations and/or obtain new ones. Like Lipid Annotator, MS-DIAL can build a fragmentation-based MS/MS library comprising the *m/z* of all precursors identified as lipids by the software and the corresponding RT. The same ion species used with Lipid Annotator were used with MS-DIAL.

Finally, manual MS/MS spectral inspection was carried out using the Agilent MassHunter Qualitative software (version 10.0), comparing the fragmentation patterns of the displayed duplicates from both software. To ensure a correct annotation, the co-elution of possible adducts for the specific lipid subclass was examined(42). The final list of putatively annotated lipids was further verified and corrected with an in-house LC-MS database of human plasma lipids(46).

### Statistical analysis

To examine the effects of HIIE, the signals of the putatively annotated lipids (n= 247) were compared. The QA of the analyzed samples (one replicate) was assessed using a PCA model showing the tight clustering of the QC samples whereas the individual points represented the truly biological samples. The number per group were the 9 healthy subjects that went to sessions A and B, analyzed individually at each matched time point (n= 6) using the Wilcoxon test (performed in MATLAB) and establishing a significant p-value for paired test at p < 0.05. In addition, to control for type I error due to multiple comparisons, false discovery rate (FDR) was calculated by using the Benjamini-Hochberg correction, statistical significance was set with a corrected p-value (pFDR) < 0.2 for exploratory studies(47,48). This information has been included in each Figure at the legend.

The MetaboAnalyst online tool v6.0 (https://www.metaboanalyst.ca) was used to produce heat maps with hierarchical clustering(49). For lipid networks, the web-tool LINEX Lipid Network Explorer (version 1) was used (50). GraphPad Prism v9.5.1 (GraphPad Software, San Diego, CA, USA) was used to represent plots with stock bars and lipid trajectories.

## Acknowledgments

Authors would like to thank Santiago Angulo for his advice on statistics.

## Author contributions: CRediT

**H.G.** and **V.M.**: Conceptualization, Funding acquisition, Methodology, Supervision, and Writing – original draft. **C.B.**, **C.G-R.**, and **G.T.** Funding acquisition, Methodology, Conceptualization, and Supervision. **S.N.**, and **V. M.** Methodology, Investigation, and Writing – original draft. **T.M.** and **C.G-R.** Formal analysis, Data curation, and Investigation. **R.N.**, **A.V.**, and **A.G.** Data curation, Investigation, Methodology, and Writing – original draft. **A.V.**, and **A.G.** Supervision. **All authors**: Writing – review and editing.

## Sources of funding for the research reported in the article

This work was supported by the HORIZON-WIDERA-2021 Project 101079370, BiACEM: BiOMIC_AUTh, Center of Excellence in Metabolomics research.

This work was supported by the Ministry of Science and Innovation of Spain (MCIN) MCIN/AEI/10.13039/501100011033, by ERDF A way of making Europe, grant number PID2021-122490NB-I00. A.V. acknowledges financial support from “Ayudas Puente curso 24/25” from San Pablo CEU University and grant number PEJ-2023-AI/SAL-GL-27622 from the Community of Madrid for hiring a research assistant.

## Availability of data and materials

All data generated and analyzed during this study are included in this article and its supplementary information files (Supplementary Tables). The raw files of lipidomics generated in this study have been deposited in the Metabolomics Workbench under Study ID ST003807 [http://dx.doi.org/10.21228/M8VR88].

