## Supplementary information for "A lipidomic exploration of the effects of high-intensity interval exercise in healthy men after metformin intake"

Coral Barbas

CEMBIO, Centro de Metabolómica y Bioanálisis, Facultad de Farmacia, Universidad San Pablo CEU. Avda. Montepríncipe s/n, 28668 Boadilla del Monte, Madrid, SPAIN.

### Content:

### Supplementary figures:

**Figure S1**: Blood samples at same time points that were used for the univariate comparison between Exercise and Resting Sessions.

**Figure S2**: Unsupervised multivariate models (PCA) using the chemical features after QA for (**A**) all samples and QCs in ESI+, (**B**) all samples and QCs in ESI–, (**C**) only annotated lipids. **D-F**, matched PCA models of all samples without QCs.

**Figure S3:** Lipid network analysis of the statistically significant lipids between resting and exercise session at **A)** T0 (0 h) just after taking metformin and **B)** at T4 (3:00 h) 1 hour after finishing HIIE.

**Figure S4:** Lipid trajectories in exercise (in red) and resting (in blue) sessions from T0 (0 h) to T5 (6 h) for additional lipids with different trajectories compared to the lipid species with the highest fold changes; PC(16:0_18:0), LPE(22:6), SM(d40:1), Cer(d18:1/22:0), TG(56:2) and TG(42:0).

**Figure S5:** Lipid trajectories in exercise (in red) and resting (in blue) sessions from T0 (0 h) to T9 (12 h) for additional lipids with different trajectories compared to the lipid species with the highest fold changes; PC(16:0_18:0), LPE(22:6), SM(d40:1), Cer(d18:1/22:0), TG(56:2) and TG(42:0).

**Figure S6**: Lipid trajectory for A) acetyl-carnitine (CAR(2:0)) and B) TG(56:2) both including the metformin concentration in exercise (in dark and light red, respectively) and resting (in dark and light blue, respectively) sessions over 24h.

**Figure S7**: **A)** Workflow of plasma sample treatment for lipidomic analysis. **B)** Workflow of lipid annotation. Icons were created using biorender.com.

### Supplementary tables (included in a separate excel file):

**Table S1:** List of the 247 annotated lipids after QA that were used for the comparison between the time points in session A and B.

**Table S2:** List of lipids that were significantly different between session A and B at T0 (0:00h). Significant differences were found using Wilcoxon test.

**Table S3:** List of lipids that were significantly different between session A and B at T3 (2:05h). Significant differences were found using Wilcoxon test.

**Table S4:** List of lipids that were significantly different between session A and B at T4 (3:00h). Significant differences were found using Wilcoxon test.

**Table S5:** List of lipids that were significantly different between session A and B at T5 (6:00h). Significant differences were found using Wilcoxon test.

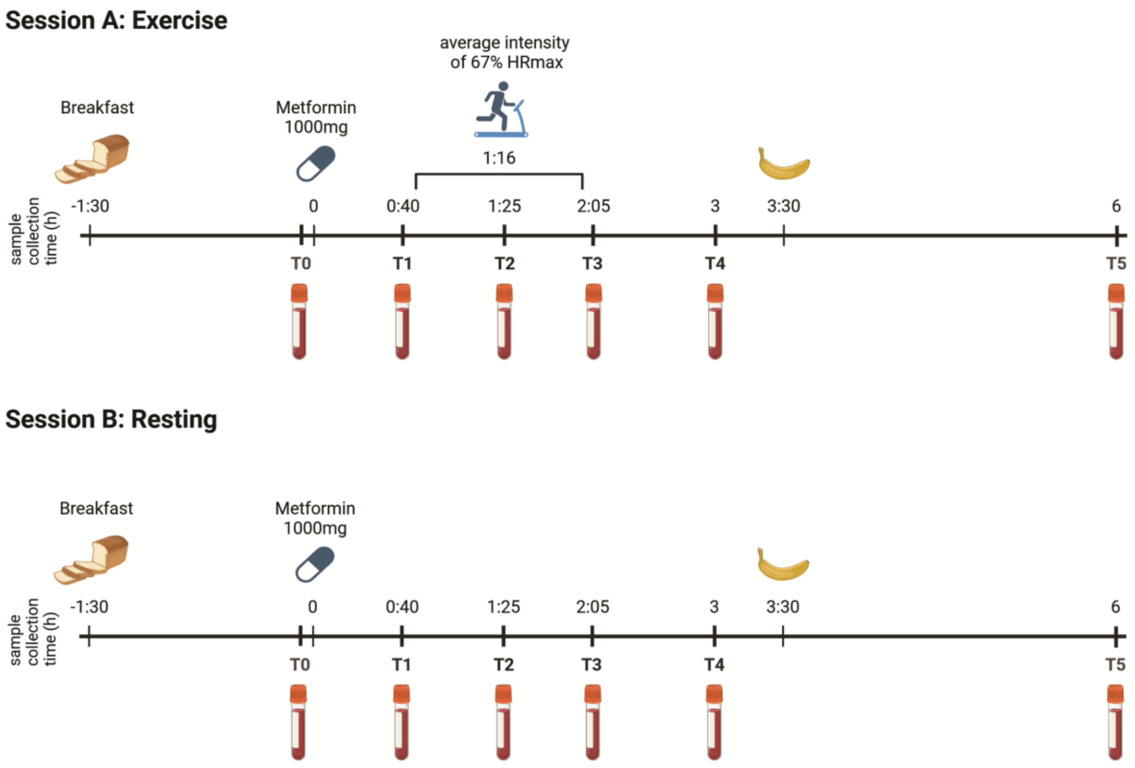


**Figure S1:** Blood samples at same time points that were used for the univariate comparison between Exercise and Resting Sessions.

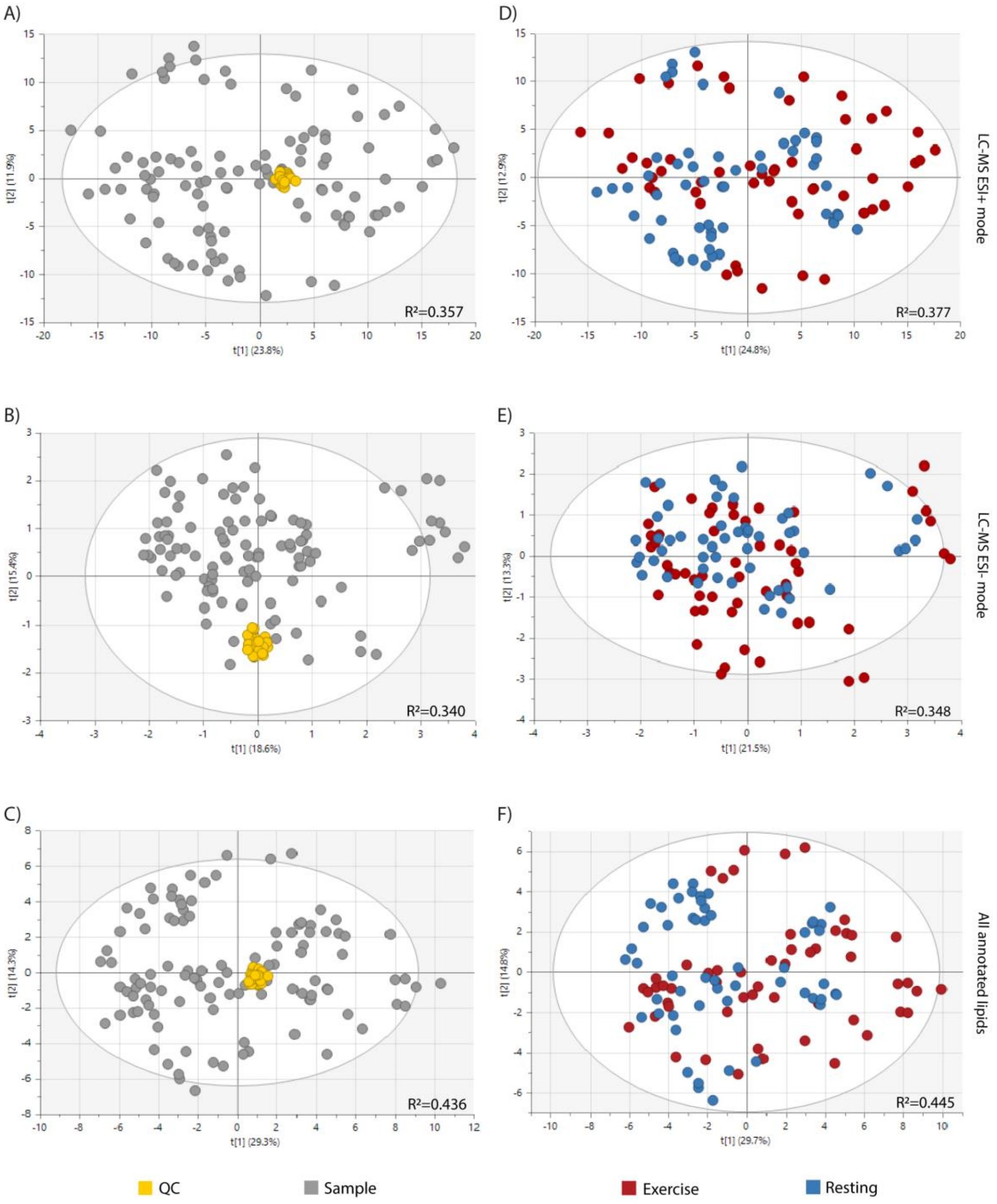


**Figure S2**: Unsupervised multivariate models (PCA) using the chemical features after QA for (**A**) all samples and QCs in ESI+, (**B**) all samples and QCs in ESI–, (**C**) only annotated lipids. **D-F**, matched PCA models of all samples without QCs, exercise in red and resting in blue. $R^2$: percentage of the variability that each component is able to explain. Hotelling ellipses depict a 95% confidence interval. Models in ESI- (B, E) were built with log transformation and centroid scaling of the data, models in ESI+ (A, D) and the merged annotated models (C, F) were built with log transformation and pareto scaling.

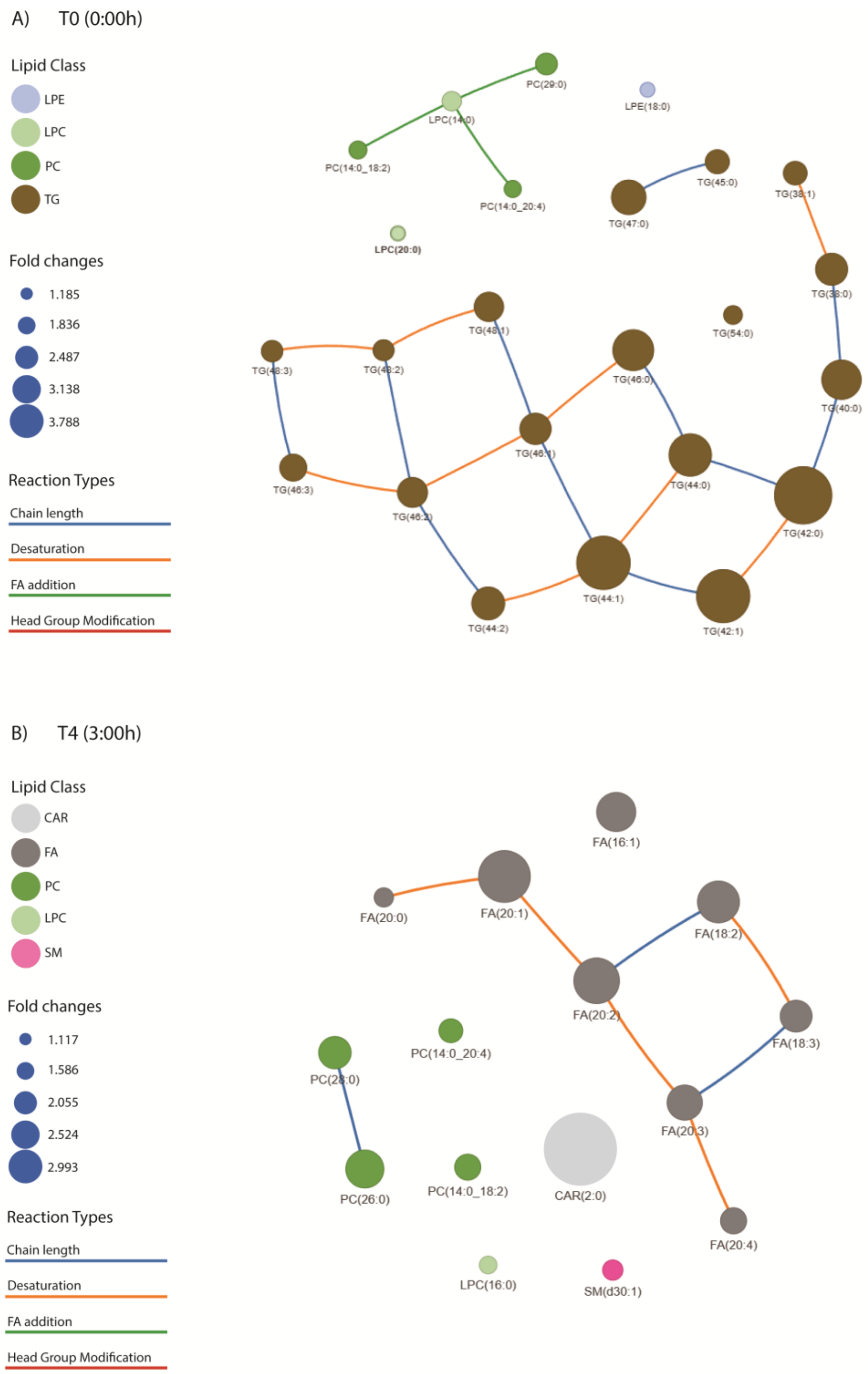


**Figure S3:** Lipid network analysis of the statistically significant lipids between resting and exercise session at **A)** T0 (0 h) just after taking metformin and **B)** at T4 (3:00 h) 1 hour after finishing HIIE. The size of the dot represents the fold change for individuals at exercise session against resting conditions. The color of the dots represents the lipid class. All lipid names are depicted as the composition ("_"), to see the sn-positions ("/") check TableS2 and S4. Lipids with the same sum composition are represented as one lipid. Abbreviations: acyl carnitines (CAR), fatty acids (FA), lysoglycerophosphocholines (LPC), glycerol-phosphocholines (PC), ether-glycerophosphocholines (PC-O/P), lysoglycerophospho-ethanolamines (LPE), sphingomyelins (SM), ceramides (Cer), hexosylceramides (HexCer), cholesteryl esters (CE), triglycerides (TG).

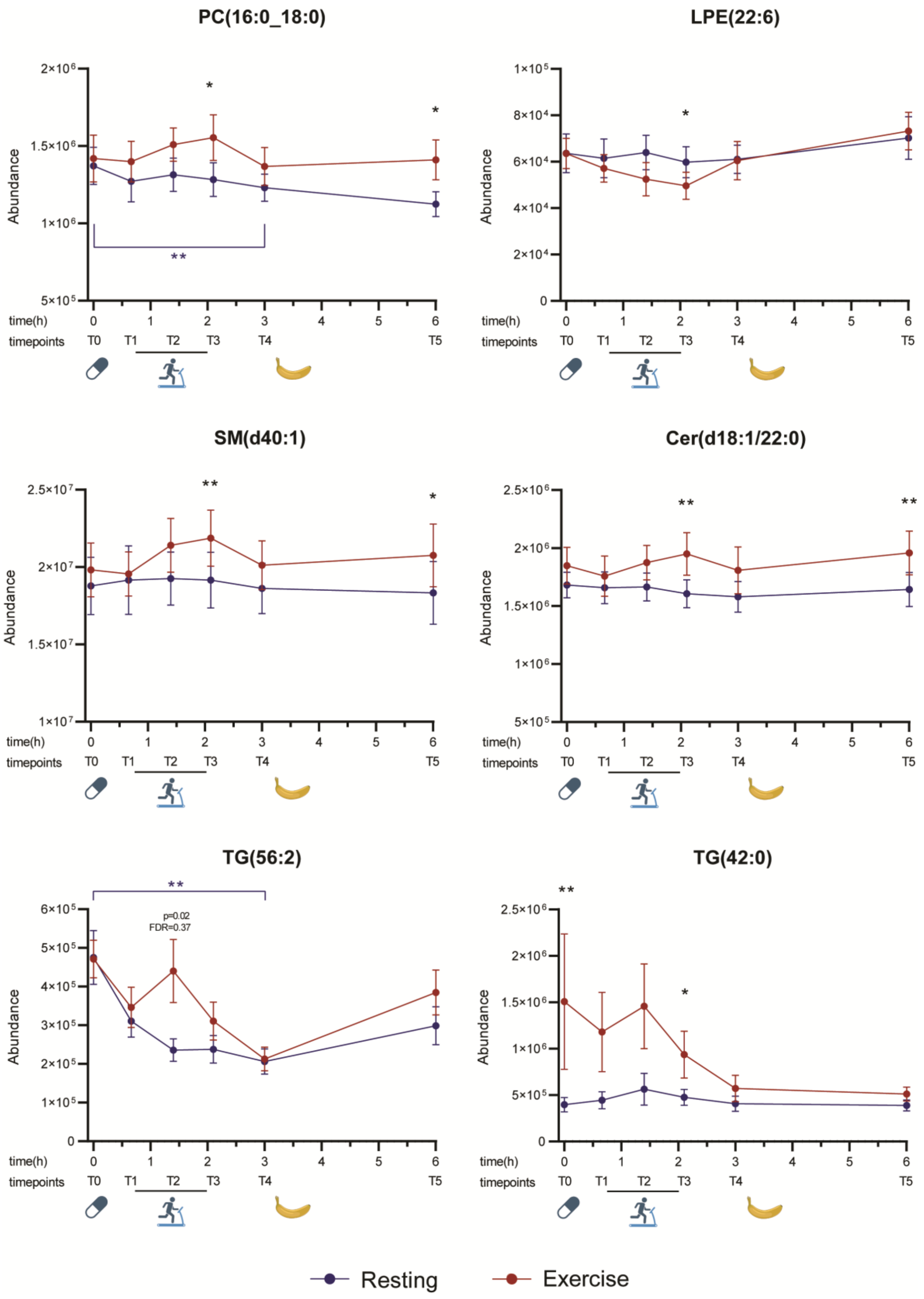


**Figure S4:** Lipid trajectories in exercise (in red) and resting (in blue) sessions from T0 (0 h) to T5 (6 h) for additional lipids with different trajectories compared to the lipid species with the highest fold changes; PC(16:0_18:0), LPE(22:6), SM(d40:1), Cer(d18:1/22:0), TG(56:2) and TG(42:0). Abundances values are represented as mean of normalized area ± SEM, significant differences were found using Wilcoxon test; * $p < 0.05$; ** $p < 0.01$.

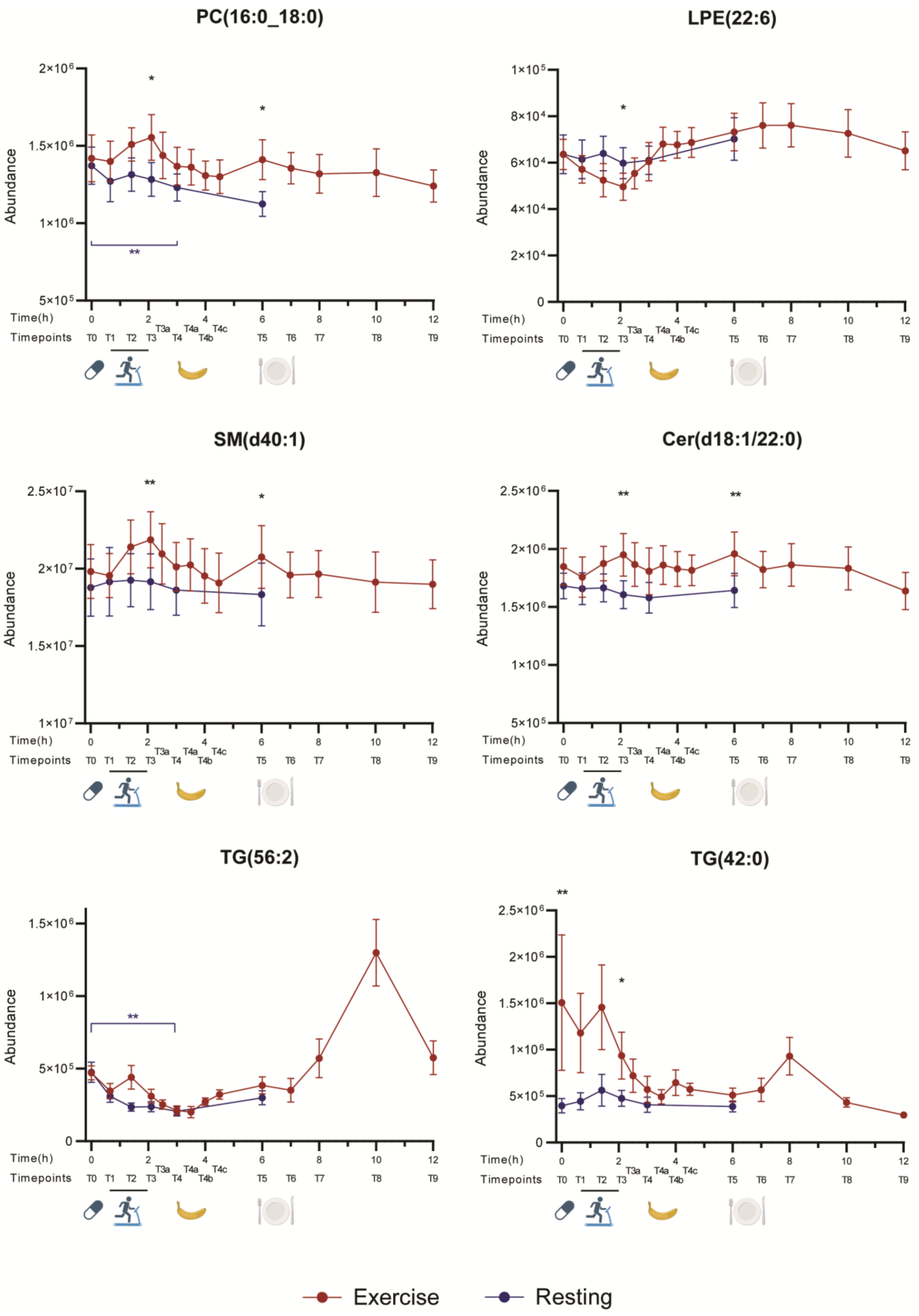


**Figure S5:** Lipid trajectories in exercise (in red) and resting (in blue) sessions from T0 (0 h) to T9 (12 h) for additional lipids with different trajectories compared to the lipid species with the highest fold changes; PC(16:0_18:0), LPE(22:6), SM(d40:1), Cer(d18:1/22:0), TG(56:2) and TG(42:0). Abundances values are represented as mean of normalized area ± SEM, significant differences were found using Wilcoxon test; * $p < 0.05$; ** $p < 0.01$.

A)

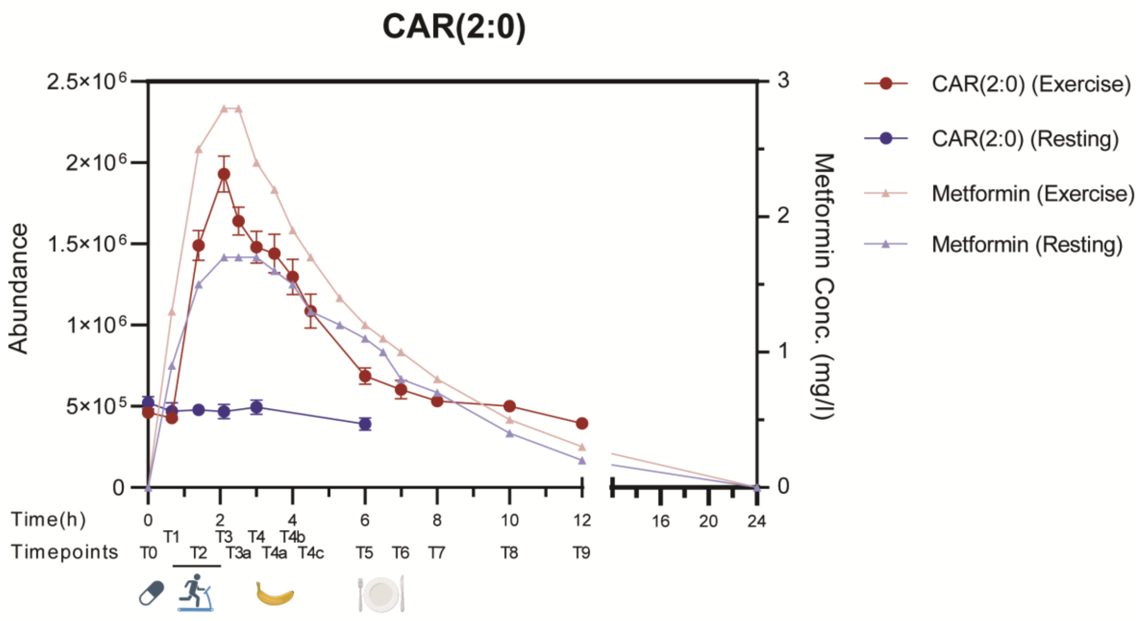


B)

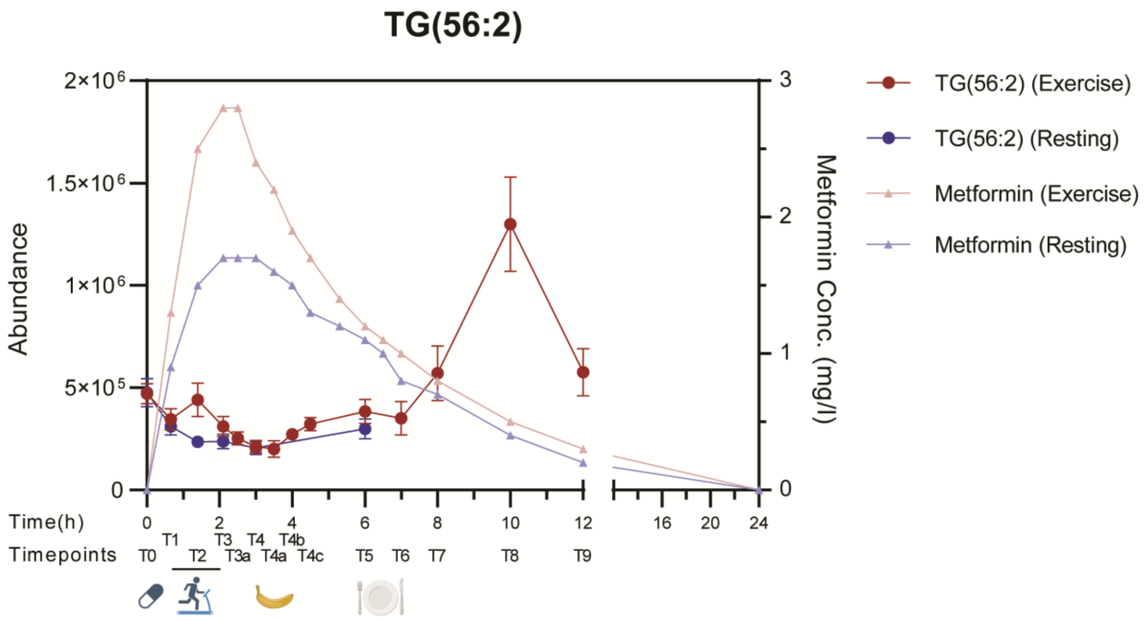


**Figure S6**: Lipid trajectory for A) acetyl-carnitine (CAR(2:0)) and B) TG(56:2) both including the metformin concentration in exercise (in dark and light red, respectively) and resting (in dark and light blue, respectively) sessions over 24h.

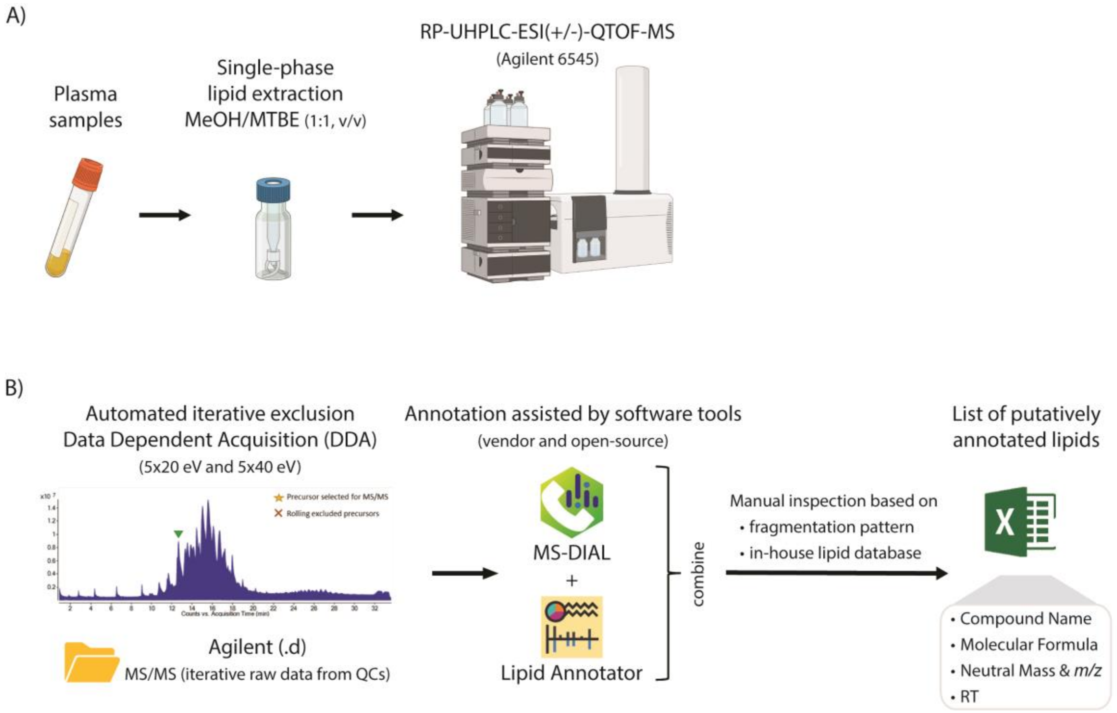


**Figure S7**: **A)** Workflow of plasma sample treatment for lipidomic analysis. **B)** Workflow of lipid annotation. Icons were created using biorender.com. Abbreviations: reverse phase–ultra high-performance liquid chromatography coupled to mass spectrometry with an electrospray ionization source and a quadrupole–time of flight mass analyzer (RP-UHPLC-ESI-QTOF-MS).
